# Quasi-neutral molecular evolution — When positive and negative selection cancel out

**DOI:** 10.1101/330811

**Authors:** Bingjie Chen, Zongkun Shi, Qingjian Chen, Darryl Shibata, Haijun Wen, Chung-I Wu

**Affiliations:** State Key Laboratory of Biocontrol, School of Life Sciences, Sun Yat-sen University, Guangzhou 510275, China; Department of Pathology, Keck School of Medicine of the University of Southern California, Los Angeles, California, USA; CAS Key Laboratory of Genomic and Precision Medicine, Beijing Institute of Genomics, Chinese Academy of Sciences, Beijing 100101, China; Department of Ecology and Evolution, University of Chicago, Chicago, Illinois 60637, USA

**Keywords:** positive selection, negative selection, cancer evolution, cancer risk, genetic drift

## Abstract

In the absence of both positive and negative selection, DNA sequences evolve at the neutral rate, R = 1. Due to the prevalence of negative selection, R∼1 is rarely achieved in organismal evolution. However, when R ∼ 1 is observed, it does not necessarily indicate neutral evolution because positive and negative selection could be equally strong but in opposite directions - hereby referred to as quasi-neutrality. We now show that somatic-cell evolution could be the paradigm of quasi-neutral evolution for these reasons: 1) Quasi-neutrality is much more likely in small populations (size N < 50) than in large ones; 2) Stem cell population sizes in single niches of normal tissues, from which tumors likely emerges, have small N’s (usually < 50); 3) the genome-wide evolutionary rate across tissue types is close to R = 1; 4) Relative to the average of R ∼ 1, many genes evolve at a much higher or lower rate, thus hinting both positive and negative selection; 5) When N < 50, selection efficacy decreases rapidly as N decreases even when the selection intensity stays constant; 6) Notably, N is smaller in the small intestine (SmI) than in the colon (CO); hence, the ∼ 70 fold higher rate of phenotypic evolution (observed as cancer risk) in the latter can be explained by the greater efficacy of selection, which then leads to the fixation of more advantageous mutations and fewer deleterious ones in the CO. Under quasi- neutrality, positive and negative selection can be measured in the same system as the two forces are simultaneously present or absent.

## Introduction

The effects of positive and negative selection are jointly reflected in the evolutionary rate of DNA sequences of coding regions. However, the landscape of molecular evolution is dominated by neutral variants as well as deleterious mutations yet to be eliminated, which collectively underpin the neutral theory (1, 2). Hence, positive selection, swamped by negative selection, is difficult to determine (1-4).

To incorporate both positive and negative selection, we let R be the rate of sequence evolution, relative to the neutral rate. The value of R can be expressed as the Ka/Ks ratio, where Ka and Ks are, respectively, the number of nonsynonymous (or amino acid-altering) and synonymous mutations per site between two DNA sequences. R is thus the net outcome between positive and negative selection that speeds up and slows down nucleotide substitutions, respectively by a fraction A and B. Hence, R = 1 + A – B when both are in action. The challenge is to tease apart the opposing factors of A and B, given R. In the evolution between species, the genome-wide average of R (designated R*) is usually < 0.3 while R* > 0.5 has not been reported in organismal evolution. When R < 1, negative selection is obviously in action but positive selection may or may not be operative.

For a few genes in the genome, R approaches 1 and the conventional view posits very weak selection such that A ∼ 0 and B ∼ 0.Note that the conventional view includes the nearly-neutral model (5). Importantly, while the conventional view of neutrality predicts A ∼ 0 ∼ B, the condition of R ∼ 1 in fact permits A ≫ 0 and B≫ 0 as long as A ∼ B. In such a system, positive selection and negative selection are both strong, but equally so in opposite directions. We shall refer to such systems of R ∼ 1 as “quasi-neutrality”, in contrast with the conventional “functional neutrality”. The issue is how often, and under what conditions, would the two opposing forces cancel each other out.

In this context, the evolution in somatic tissues that eventually transitions to tumorigenesis is relevant (6-11). This process has ultra-microevolutionary distances, in the order of 10^−5^-10^−6^ per bp, far smaller than the evolutionary distances between closely related species, which are typically 10^−2^ - 10^−1^ per bp. Properties at the “ultramicro-evolutionary” scale are often distinct from those in the evolution between species (8). In this study, we ask whether somatic evolution could be quasi-neutral evolution in action. Systems of quasi-neutrality will be of general significance in molecular evolution because the detection of positive selection in organismal evolution is uncertain. These analyses usually rely on additional data and make further assumptions (12, 13) but such assumptions may often be mutually incompatible (He *et al.* Unpublished). The ability to analyze positive and negative selection in the same genome may therefore shed new light on the concurrent operation of the two opposing forces.

## RESULTS

### I On the joint influences of positive and negative selection - Theory

With both positive and negative selection in operation, the rate of molecular evolution can be expressed as

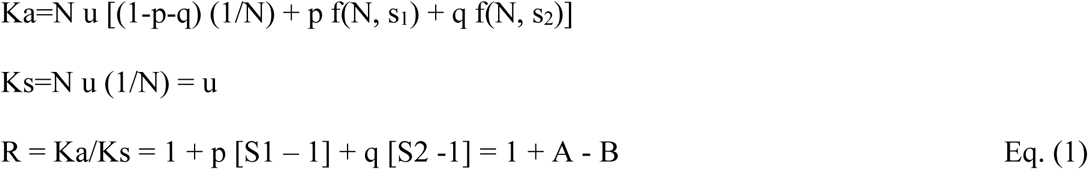

where Ka is the number of nonsynonymous substitutions per nonsynonymous site and Ks is the corresponding number for synonymous changes. In Eq. (1), A = p [S1 – 1] and B = - q [S2-1], where p and q are the proportion of mutations under positive and negative selection, respectively. S1 and S2 are defined as the rate of accumulating fitness-altering mutations relative to the speed of neutral evolution. The theory has been well described in textbooks (2-4):

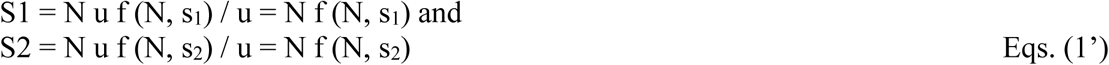

In Eq. (1’), f (N, s) = (1-e^-2s^)/ (1-e^-2Ns^) is the fixation probability of non-neutral mutations where s is the selective coefficient with s_1_>0 being beneficial and s_2_<0 being deleterious.

Fig. 1A shows how the efficacy of selection (S1 and S2) changes, in the log-scale, for both s>0 and s<0 when N varies between 1 and 20. Fig. 1B provides finer details in the linear scale for s>0. Note that the efficacy increases as N increases and the trend is much stronger for negative selection than for positive selection. Here, we set s_1_ = -s_2_ = s but more complicated setups (14) would yield the same qualitative conclusion. Assuming s < 1 and 2Ns > 1, Eq. (1) can be simplified as:

**Fig. 1.**
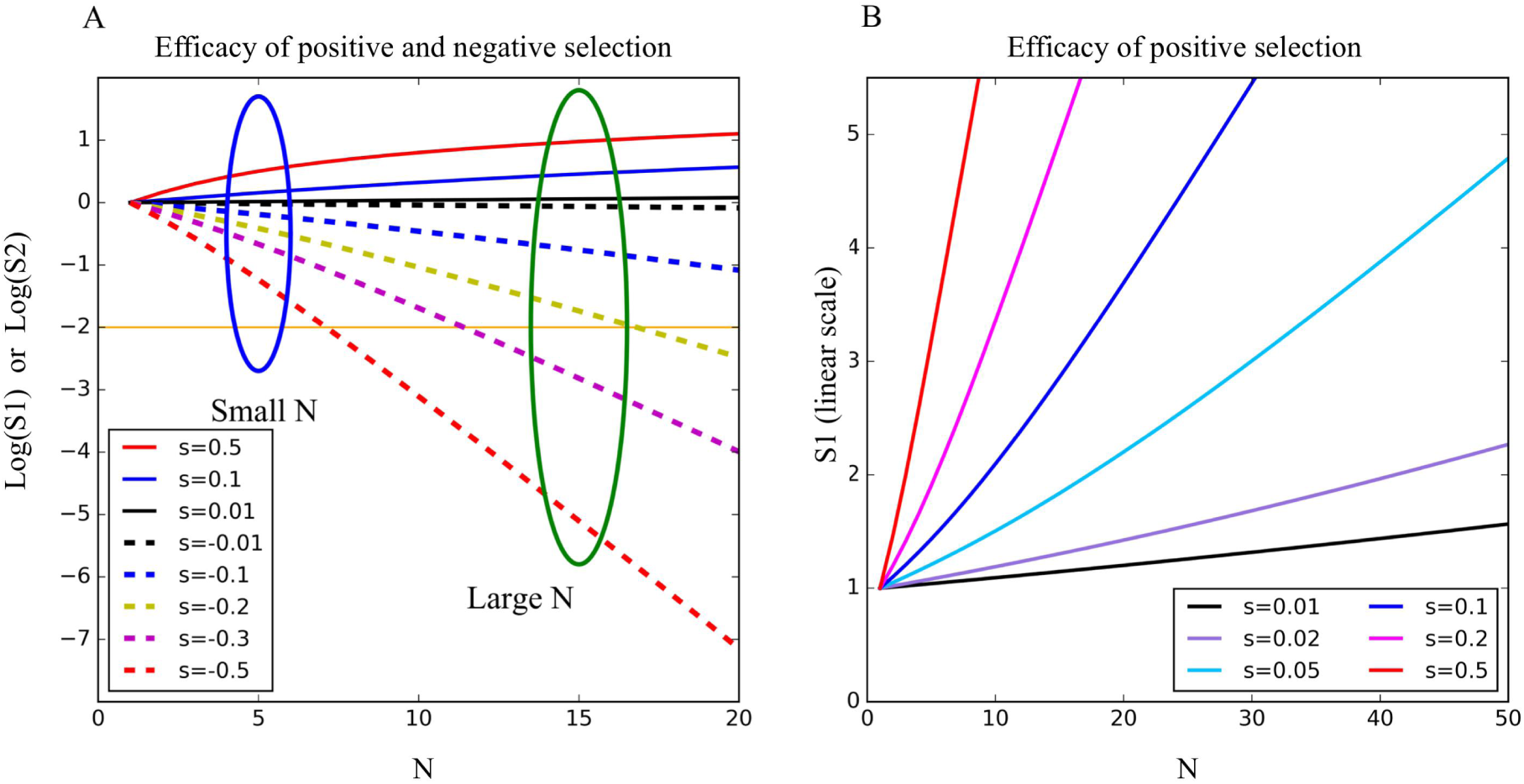
The rate of molecular evolution as a function of N. (A) Log (S1) and Log(S2) (solid and dashed lines, respectively) on the Y axis designate the efficiency of selection (Eq. 1’). It represents the rate of accumulating advantageous or deleterious mutations relative to the neutral rate, as a function of N (population size). The orange horizontal line indicates S2 = 0.01 above which fixation is detectable. The speed of fixing mutations in small vs. large population is contrasted by the blue vs. green oval. (B) A higher resolution plot for advantageous mutations. The scale of the Y-axis is in the linear scale.

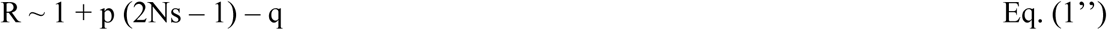

The effect of positive selection, relative to that of negative selection, is amplified by a factor of (2Ns -1) in Eq. (1’’). When the two forces are in near parity, they yield A ∼ B and R ∼ 1. Hence, p = 0.1, q = 0.9 and 2Ns = 10 is a quasi-neutral system even though all mutations are under selection. Such a “quasi-neutral” system does not require A ∼ 0 ∼ B, as does the conventional neutrality.

Fig. 2 illustrates the R values for 2Ns = 2, 5, 20 and 50 under different proportions of (p, q) using Eq. (1). We highlight the range of (p, q) that satisfies quasi-neutrality with Ka/Ks between 0.85 and 1.15 (shaded red). When Ns is small, the range is quite broad but decreases very rapidly as Ns increases. For example, when 2Ns = 50, p has to be smaller than 0.02 and q has to be large. Notably, a small change in p is accompanied by a large jump in q. The analysis shows that quasi-neutrality is increasingly likely when Ns becomes smaller because A→ 0 and B→ 0 and therefore A∼B is more likely.

**Fig. 2.**
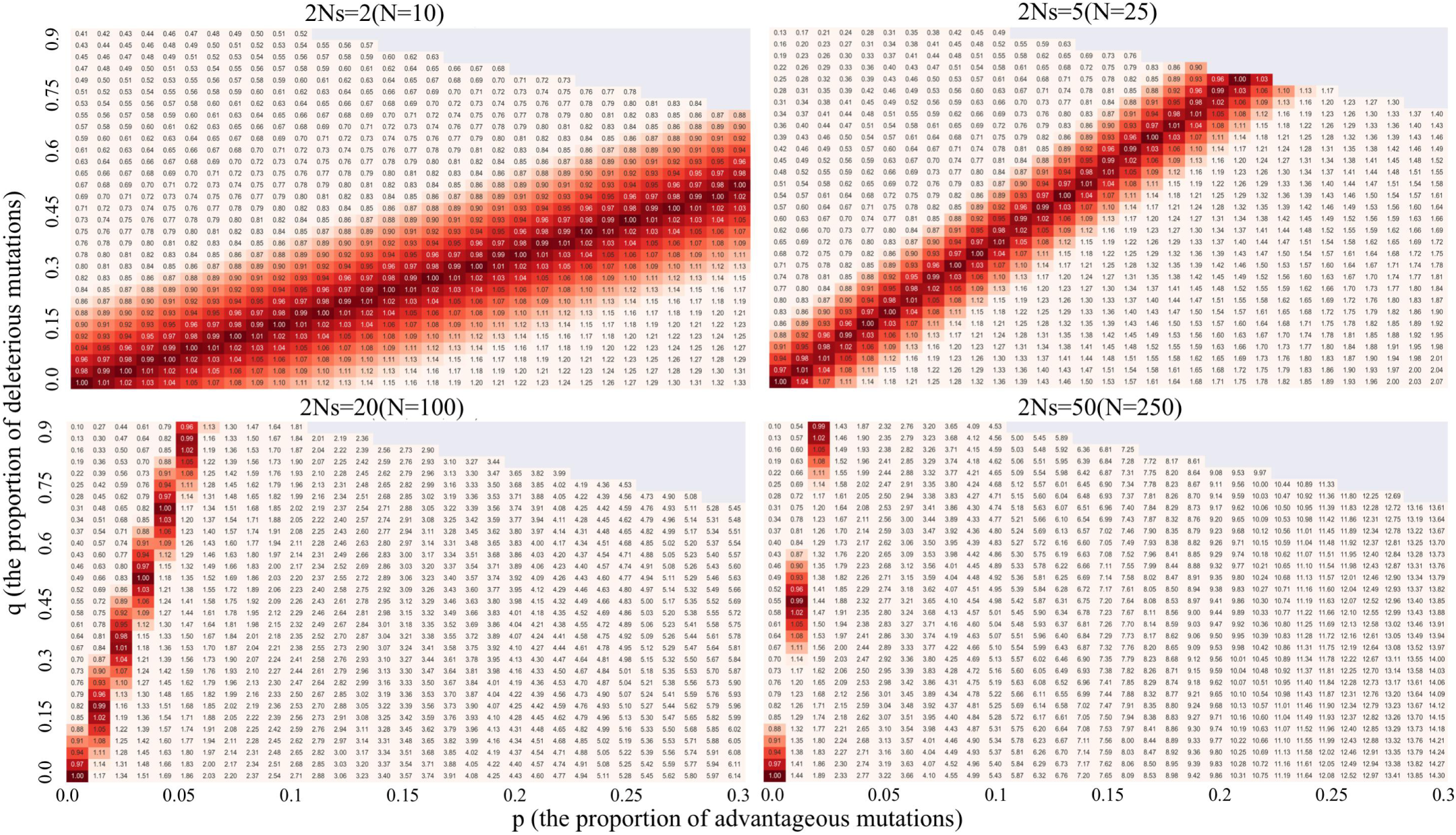
Heatmap of R (Ka/Ks) ratios according to Eq. (1) under different combinations of p’s and q’s on the X and Y axis. Respectively, p and q designate the proportion of advantageous and deleterious mutations. Shaded area portrays R values in the range of 0.85-1.15 centered around R∼1 (the darkest shade). The four panels represent 2Ns = 2, 5, 20 and 50 while s = s1 = -s2 = 0.1. Note that the shaded area is broad when 2Ns=2 but decreases rapidly as Ns increases. Thus, quasi-neutrality is more likely as Ns gets smaller.

### II Somatic-cell evolution as the paradigm of quasi-neutral evolution

A likely system of quasi-neutrality is somatic-cell evolution, which could lead to tumorigenesis at times. The stages of somatic-cell evolution are depicted in Fig. 3a (see ref. (8) for detail). We shall focus on Stage I, which ends in the single cell labeled c-MRCA (the most recent common ancestor of all cancerous cells). The c-MRCA cell demarcates Stage I and II, right at the very beginning of tumor growth. The evolution in Stage I most likely takes place in the somatic stem cell niches. As depicted, few cells evolve to become c-MRCA. Hence, the comparisons between cells that do and do not reach c-MRCA will be crucial in later sections.

**Fig. 3.**
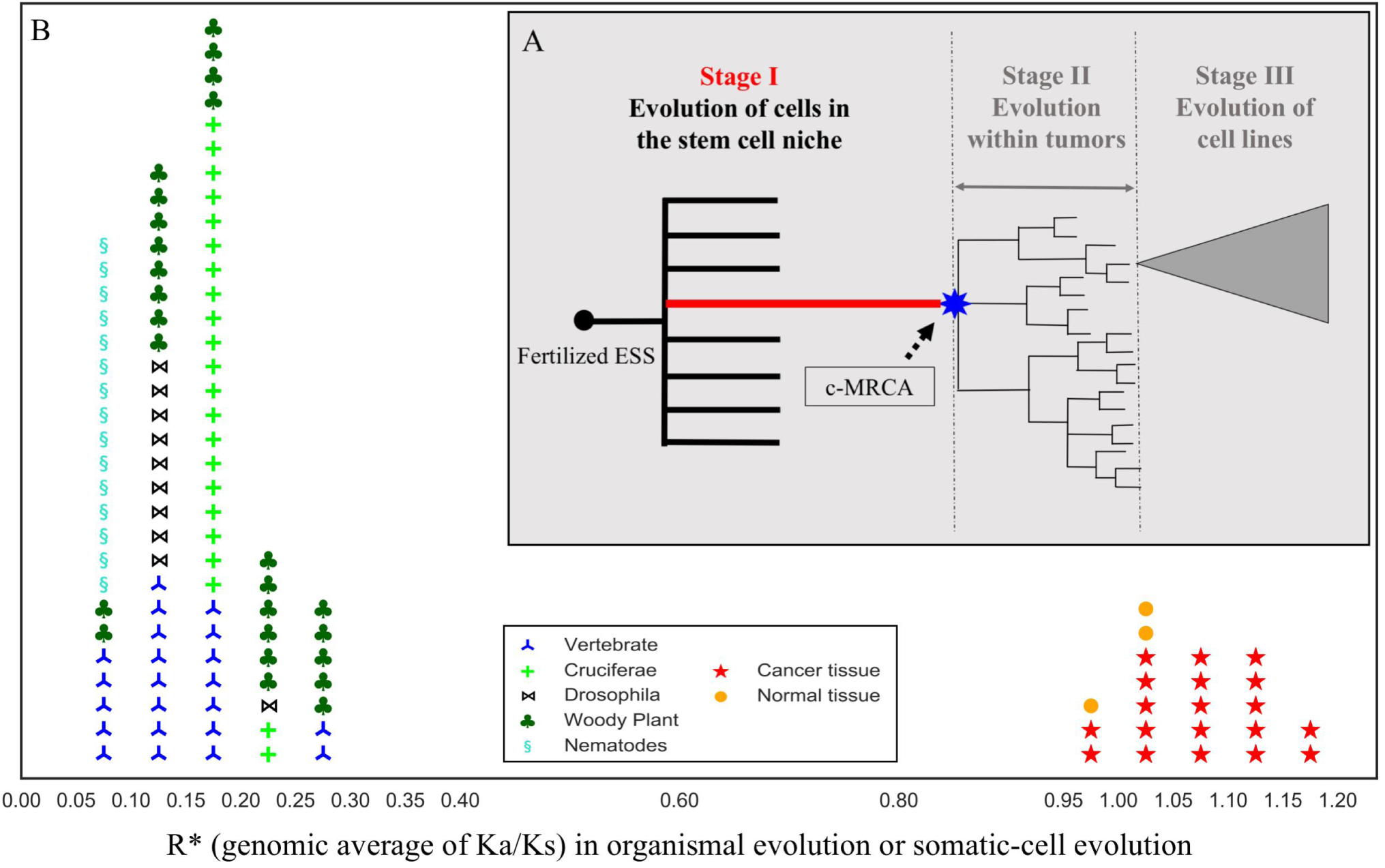
(A, inset) Stages of somatic cell evolution. Stage I, starting in a stem cell and ending in the c-MRCA (the most recent common ancestor of all cancer cells of a tumor), is the focus of this study. DNA sequences of tumors (9, 10) reflect the evolution in this stage. Stage II represents the diversification within tumors (22, 43, 45, 46) and Stage III is the further evolution in cell lines (47). (B) The distributions of R* (genome-wide Ka/Ks ratio) for natural species and for somatic tissues (both normal and cancerous samples) are shown. Sequenced genomes of natural species from a wide range of taxa (vertebrates, insects, nematodes and plants) generally have an R* ratio between 0.05 and 0.3 (clustered on the left). In contrast, across 19 cancer types and 3 normal tissues (21), the R* values are clustered between 0.95 and 1.15. The tissue types are given in the Supporting Information.

Somatic mutations reported in tumor sequencing projects (9, 10) represent the genetic make-up of the c-MRCA cell, or Stage I evolution (see Fig. 3a and legends). Fig. 3b shows that, across 22 tissue types, the estimated R*’s are close to 1, ranging between 0.97 and 1.20 with a grand mean of 1.08. In comparison, sequenced genomes of natural species generally have an R* ratio between 0.05 and 0.3. Because the evolution of somatic cells differs from that of organisms in many respects, notably the absence of recombination, the population genetic theory has often been questioned (see ref. (8)). The supporting information addresses the issue, noting that recombination does not affect the mean evolutionary rate (e.g., Eq. 1 -1”). Recombination does affect the variance, but only when the rate itself is high.

While Fig. 3b portrays somatic cell evolution as compatible with the quasi-neutrality of Fig. 2, it could also be compatible with the strictly neutrality of p ∼ q ∼ 0. Indeed, a recent report suggested q ∼ 0 in somatic cell evolution although, curiously, it also estimated p to fall between 1% and 4% (15). Earlier, p was suggested to be ∼5% (8). According to Eq. 1’’, R ∼ 1 dictates q > p if p > 0. Even if Ns is as small as 3, merely three times as large as the neutrality of Ns =1, q would still be ∼ 5p. A companion analysis (Chen QJ *et al*.; see Appendix) provides further justifications for q > p > 0 to account for the TCGA data. In short, somatic cell evolution is not likely to be neutral in the conventional sense.

#### Size of stem cell population in a niche

Fig. 2 shows that 2Ns must be small to have R* close to 1. Thus, N and s cannot both be large. In particular, if N is large (> 100), s has to be small and the system reverts to the conventional neutrality. Here, we evaluate the range of N in somatic evolution. As portrayed in Fig. 3a, the relevant stage happens in the stem cell niche, from which tumors likely emerge. Each individual niche is the unit of evolution, within which competitions between cells happen (6, 16-21). Critically, it is the size of the stem cell population in a niche (N) that determines the efficacy of selection.

A stem cell niche in the small intestine (SmI) and colon (CO) is in the crypt (17, 18) where N is generally suggested to be in the range of 2 – 20 in mouse, and comparable in human (19, 20). Since a crypt in the human SmI is roughly 1/5 as large as a colonic crypt (Fig S1), it is hypothesized that N in the SmI (N_SmI_) is smaller than in the colon (N_CO_). To test the hypothesis of N_SmI_ < N_CO_, we cultured organoids from single crypts collected from human SmI and CO (see Methods). Simulations suggest that genome sequencing data from single crypts can be useful for this test.

Briefly, sequencing data from a sample of n cell lineages would yield a small peak for low frequency variants (Fig. S3) at the interval of 1/2n where 2n is the total number of genomes in n cells. Such a peak would be detectable if n is between 2 and 10. In our data, sequenced SmI crypt samples reveal a visible small peak that suggests N_SmI_ to be in the range of 2 – 5 (see supporting information. The peak is missing in the CO data, thus suggesting N_CO_ > 10, which has been shown to be < 30 (19, 20). Given the range where N_SmI_ and N_CO_ fall, the evolution in somatic tissues would satisfy the quasi-neutrality condition of R ∼ 1 even when s is not small (say s > 0.1).

### III Testing the quasi-neutral model in Stage I of somatic-cell evolution

Given 50 > N_CO_ > N_SmI_ > 2, the two predictions under quasi-neutrality are:

i. At the DNA sequence level, CO is predicted to have accumulated more advantageous mutations but fewer deleterious ones in comparison with SmI, resulting in comparable R*’s (∼1) between the two tissues.
ii. At the phenotypic level, CO would evolve the proliferative phenotype at a higher rate, and hence would have a higher rate of phenotypic evolution (observed as cancer risk), than SmI.

#### Rate of DNA sequence evolution

For the first test, we separate nonsynonymous mutations into advantageous, neutral and deleterious ones following the procedure of Wu *et al*. (8). In a collection of 12,420 genes of colon adenocarcinoma (COAD; from TCGA open portal), 3.6% of them, with Ka/Ks > 4, are classified as advantageous and 23.8% of the genes that show a Ka/Ks ratio of < 0.5 are classified as deleterious. The rest are considered neutral. Here, we compare the distributions of non-synonymous changes in the three categories from the normal SmI and CO (21) as well as from the COAD (TCGA open portal). Note that q/p > 1 (at 6.1 = 23.8%/3.6%) is as expected in Eq. (1’’).

In the normal SmI, the distribution of SNVs is very close to the distribution of the gene number itself (Table 1, 4.3%-73.5%-22.2% vs. 3.6%-72.7%-23.8%, *P* = 0.88 by G-test) as if SNVs simply fall randomly on the genes with little interference by natural selection. The normal SmI tissue indeed appears to evolve neutrally. On the other hand, the distribution of SNVs in the normal CO is distinct from the gene number distribution (5.0%-78.6%-16.4% vs. 3.6%-72.7%-23.8%, *P* < 0.01 by G-test), suggesting that these mutations do not come randomly from all genes. Positively selected genes tend to contribute more SNVs in the normal CO and negatively selected genes contribute fewer than expected. SNV distributions in the colorectal adenocarcinoma tissue (COAD, 11.6%-82.3%-6.1%) strongly deviate from those of the two normal tissues by having far more advantageous mutations and far fewer deleterious ones (*P* < 0.001 by G-test).

**Table 1.**
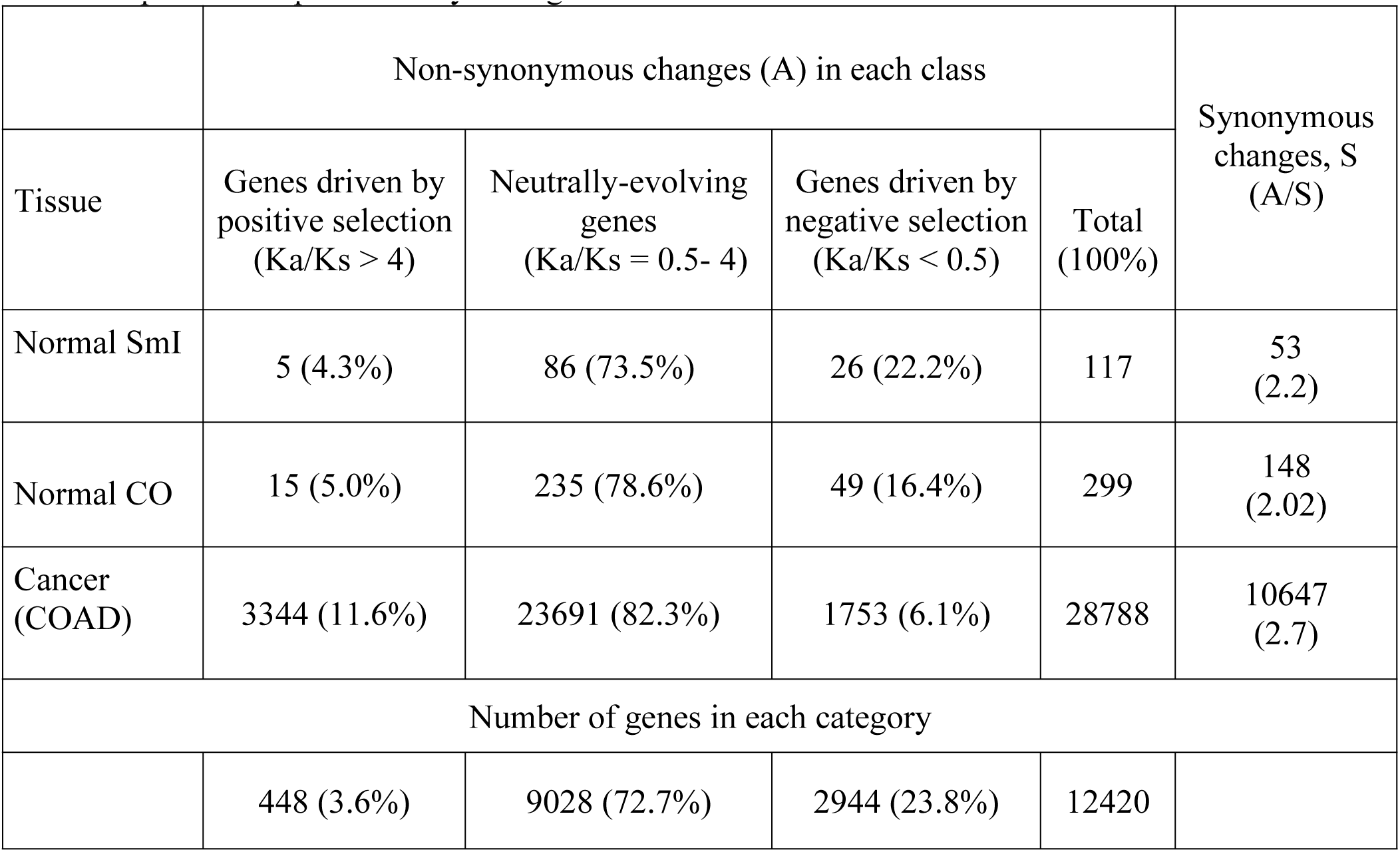
Distributions of nonsynonymous changes (A) in the 3 classes of genes, delineated by the overall selective pressure experienced by each gene in the tumors.

The distribution of SNVs progressively deviates from the random pattern going down Table 1, as positive selection becomes more potent in fixing good mutations and negative selection becomes more effective in eliminating bad ones. This is also the pattern portrayed in Fig. 1. We should also note that SNVs of each gene in Table 1 are a mixture of all three types of sites, but with biases toward one type, while Eq. (1’) models pure types of sites.

#### Rate of phenotypic evolution (i.e., cancer risk)

Between the CO and SmI, the ratios of advantageous/deleterious mutations are different. One therefore expects the crypts in the CO to evolve the proliferative phenotype more speedily. Indeed, the evolution rate (or cancer risk) in the CO is more than 70-fold higher than in the SmI (https://seer.cancer.gov). This difference is another line of evidence supporting quasi-neutral cancer evolution as strictly neutral evolution, by definition, should not have a fitness-altering consequence. The two predictions may potentially help explain how normal CO and SI accumulate similar numbers of age-related mutations (25) but have very different cancer rates (see below).

### IV Strength of selection under quasi-neutrality

It is now feasible to estimate the strength of selection. By considering both sequence evolution and phenotypic evolution, our estimation is very different from previous attempts (8, 11, 22, 23).

#### Efficacy of positive selection as a function of N

The strength of selection (S1 of Eq. (1’)) is the rate of accumulating advantageous mutations relative to the neutral rate. S1 can also be expressed as the function of the cancer risk (C_R_ = the proportion of the population with the cancer) and the R value (see Methods for details). Hence,

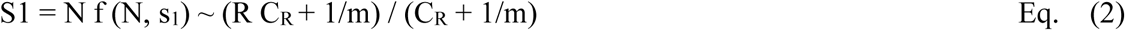

where m is a relative measure of K_S_ in cancer vs. normal tissues, usually falling between 3 and 5. Both R and m can be obtained from the TCGA data and studies of normal tissues (see Methods).

For the fastest evolving gene, P53, we use R ∼ 152, m∼4 and C_R_ = 0.048 (the latter from https://seer.cancer.gov) to obtain S1 = 25 for colorectal tumors. Substituting S1 = 25 into Eq. (1’), we obtain s_1_= 0.896 and 0.347 when N is 30 and 50, respectively. (When N is smaller than 30, s_1_ would be even larger). This level of selection is never attained in species evolution although some estimates of artificial selection are larger (24, 25). Given that P53 is the only gene in the genome that is mutated in more than 40% of all cancer cases, selection has been known to be exceptionally strong.

We also use the top 1% of genes with the highest R values (R > 4) to estimate s. For a gene with R=4, S1 = 1.48. Substituting this number into Eq. (1’), we obtain s_1_ = 0.048, 0.015 and 0.0086 for N = 10, 30 and 50, respectively. These s_1_ numbers are perhaps the first estimation of selection that takes into account the rate of phenotypic evolution in a stochastic framework. By these estimates, the most strongly selected 1% of genes in the normal CO experience a selective advantage of 1% - 5%. In populations with N <= 20 (as is likely in SmI), selection of this intensity is effectively neutral because s is no larger than 1/N. For that reason, the normal tissue in SmI is not expected to accumulate more advantageous mutations than random accumulations, a suggestion corroborated by Table 1.

#### Efficacy of negative selection as a function of N

We now address deleterious mutations. In general, deleterious mutations are eliminated, rather than fixed. The exception would be in small populations or species that have large non-recombining genomes. Under such circumstances, deleterious mutations would accumulate by a one-way degenerative process referred to as the “Muller’s ratchet” (26). The ratchet moves a notch whenever the population fixes a new deleterious mutation and eventually saddles the population with defects, thus reducing its fitness. Since SmI has a much smaller N in the crypt, deleterious mutations would accumulate at a faster rate there than that in the colon (S2) (see dotted lines of Fig. 1A). Thus, Muller’s ratchet moves faster in the SmI than in the CO.

Fig. 1 shows that the selection efficacy for N = 2 and N=20 is 0.99 and 0.82, respectively, when s_2_ = −0.01. The rates become 0.900 and 0.083 when s_2_ = −0.1 with an 11-fold difference. The difference increases as the effect of negative selection getting stronger. Since the normal tissues accumulate fewer than 100 coding mutations per genome (Table 2), the number on the Y-axis of Fig. 1A needs to be > 0.01 to have at least one deleterious mutation (above the horizontal orange line). Thus, in reference to Fig. 1A, selection against deleterious mutations should be |s_2_| < 0.2. At this strength, N between 15 and 20 would permit deleterious mutations to be fixed with a probability of 0.01 but, when N is around 5, the probability would be 21-fold higher. It is thus plausible that the SmI would have a much higher load of deleterious mutations than the CO, which was observed in the data (Table 1).

**Table 2.** Mutation accumulation, U (Number of mutations per Mb), in normal tissues

|  | Normal tissue |  |  |  |
| --- | --- | --- | --- | --- |
| Method | Liver | Colon | Small intestine | Average |
| Single cell cloning (Ref. 25) | 0.58<br>[0.48 – 0.68,<br>n =8] | 0.93<br>[0.78 – 1.20,<br>n=16] | 0.84<br>[0.56 – 1.28,<br>n=6] | 0.80 |
| Crypt cloning (this study) | -- | 0.77<br>[0.79, 0.87]<sup>a</sup><br>[0.63, 0.77]<sup>b</sup> | 0.64<br>[0.63, 0.65] | 0.71 |

<sup>a</sup> Ascending colon
<sup>b</sup> Distal colon

We note that “Muller’s rachet” in the crypts has been addressed previously (27, 28) but the emphasis and conclusion are quite different from those of this study. Here, we conclude that quasi-neutrality has significant phenotypic and clinical consequences. Cancer risks are very different between the colon and small intestine due to different efficacies of selection.

### V Selection or mutation driving the divergent cancer risks?

In this study, the cancer risk, C_R_, is assumed to be selection-driven as Fig. 1 depicts. However, many studies in the cancer literature used to, and still do, assume that C_R_ is mutation-driven (29, 30). In this view, the greater cancer risk in the CO than in the SmI is due to the divergent rates of somatic mutation. If that is true, C_R_ should be strongly correlated with each normal tissue’s mutation accumulation (referred to as U). Such a correlation has been claimed and disputed (30-32) in the absence of the direct measurement of U’s in somatic tissues.

Direct measurements were first reported by Blokzijl *et al* (21). As shown in Table 2, U ranges between 0.5 – 1.3 per Mb, about half of the variation could be attributed to age (50 – 80 yrs). These U estimates are not far from the median U value at 1-5/Mb in many human cancers (9, 10). This raises the possibility that U might have been over-estimated in the culturing of single stem cells. Because measuring the mutation burden of a single cell, or a single DNA molecule, requires a step of amplification, single cell measurements may be biased upwards by errors in the first few rounds of amplifications(33). Our organoid sequencing data thus provide a confirmation of single cell measurements.

The results of Table 2 show that whole-crypt mutation numbers are only 20 – 25% smaller than the numbers from single-cell culturing (P > 0.05). Note that, unless the clonality is 100% and all stem cells in a crypt are identical, the U values of whole crypts are expected to be lower than the single cell measurements. This small difference indicates that single cell culturing does not substantially inflate the mutation number, if at all. The finding that cancer risk does not depend on U has turned out to be general (21, 34-37). The conjecture of Cairn (6) that the tissue architecture reduces the long-term accumulation of mutations can be generalized: mutation accumulations across tissues converge into a small range. Thus, the variation in C_R_ across tissues is not directly attributable to the variation in mutation number.

## Discussion

Tumor progression has been accepted as a process of evolution since Nowell’s (7) and Cairn’s (6) seminal papers. Somatic-cell evolution, however, often appears to show different evolutionary patterns from organismal evolution (8). Even the basic concept of random genetic drift is perceived very differently (38). The most dramatic difference between somatic-cell and organismal evolution is, of course, their divergent evolutionary rates as shown in Fig. 3b.

The issue is whether the two evolutionary processes can be unified conceptually using the same framework but different parameter values. Alternatively, one might suggest that two distinct models are necessary. For genetic drift, a simple reformulation of the concept should work for both processes (38). For natural selection, the pattern in Fig. 3b is usually interpreted to mean distinct actions of selection; for example the near absence of negative selection in somatic-cell evolution (15) whereas it dominates organismal evolution. In this view, cancer cells would be unique complex systems that are robust in the face of random perturbations (Chen QJ *et al*.; see Appendix). Other scenarios in the literature amount to group selection, reciprocal altruism, spite behavior as well as other unusual modes of selection (8, 39-41) may entail a drastically modified framework.

In this study, we found that the effective population size (N) alone may be sufficient to explain the distinct evolutionary patterns between somatic-cell and organismal evolution under the same framework (Fig. 3). Classical population genetic theory assumes a sufficiently large N (≫ 100). In contrast, because stem cell niches usually have very small population sizes (N < 50), the proportion of mutations under positive and negative selection can be very sensitive to small variations in N. Organismal and somatic-cell evolution can thus be understood using the same framework. In this sense, somatic-cell evolution can be a new testing ground of evolutionary theories that have eluded empirical tests for want of suitable organisms.

## Methods

### Processing of tissue Samples

The patient was a 44-year-old female with no detectable cancer or other intestinal disease. Informed consent was obtained from the patient. Biopsy samples from the small intestine, ascending colon, and distal colon were isolated with an enteroscope, followed by organoid culture for 2 weeks. Single crypts were made up of fewer than 2000 cells. Crypts from the small intestine were composed of fewer cells than colon as the ratio of crypt sizes is about 1:5. (Fig. S1). DNA was extracted when the single organoid grew to around 10,000 cells (Fig. S2) using the QIAamp DNA Micro extraction kit.

### Whole-genome sequencing, alignment, point mutation calling, and filtering

We used the Ovation^®^ Ultralow System (V2 1–16, NuGEN) to prepare standard NGS libraries. The libraries were sequenced in paired-end (2 × 151 bp) runs using Illumina HiSeq X-Ten. Sequence reads were mapped to the human reference genome GRCh37 using the Burrows–Wheeler Aligner v0.7.15(42). Raw variants were called using the GATK HaplotypeCaller v3.7.0 with default settings and some additional options (see Supporting Information). Mutations called from different samples were combined into a single VCF file using GATK GenotypeGVCFs v3.7.0. A comprehensive filtering procedure was applied to obtain high quality somatic point mutations (see Supporting Information).

### Comparison of stem cell number in the CO vs. in the SmI

The data presented in Table S1 and Fig. S3 have led to the interpretation that the number of stem cells in a niche in the SmI is less than 5 and the corresponding number in the CO is great than 10. Let us first assume that the oganoid sequencing data come equally from n stem cells that are also genealogically equi-distant to one another (i.e., the genealogical tree of these n cells being a “star phylogeny”). In that case, the distribution of VAF (variant allele frequency) would have two peaks – a main peak at 0.5 for variants shared by all cells and a smaller peak at 1/2n.

Based on the data of Blokzijl et al.(21), the relative size of the two peaks should be about 20:1. (The mutation numbers shared by stem cells of the same niche is about ∼2500 while each individual stem cell has 100 – 150 private mutations.) In reality, the minor peak is broader than the main peak because the n cells may not contribute equally to the data; neither would they come from a star-phylogeny. In addition, due to sequencing errors, even a single stem cells would yield a small minor peak (e.g., see Ling et al. ref (43)). The contribution of sequencing error would be increasingly prominent when n becomes larger. When n > 10, the minor peak would consist mainly of sequencing errors.

VAF clustering of Fig. S3 was performed by SciClone (44), which uses the Bayesian beta mixture model to estimate the clonal composition in each sequencing sample. The analysis confirms a major peak at VAF∼ 0.5 and a minor peak near VAF of 0.2 in the SmI sample. For the CO samples, there is only one major peak at VAF of 0.5. Because Fig. S3 shows a minor peak between 0.15 and 0.25 in the SmI data, we roughly estimate n to be between 2 and 4. If we factor in the process of organoid culturing, it would be reasonable to suggest N_SmI_ < 5. Given the sequencing depth of > 90X (with an average of ∼ 97X), simple simulations show that the absence of a minor peak in the CO sample would suggest N_CO_ > 10. Based Table S1 and Fig. S3, the private/shared mutation numbers, delineated by VAF of 0.3, are 335/1863 for SmI1, 173/2491 in the ascending CO, and 146/2224 in the distal CO.

#### Population genetic theory in relation to somatic-cell evolution

Because the evolution of somatic cells differs from that of species in many respects, notably the absence of sexual reproduction and recombination, biologists have sometimes question the validity of the theory for understanding cancer evolution. A detailed discussion is hence presented in Supporting Information.

### Deduction from efficacy of selection to the formulation of KA/KS in cancer and cancer risk

The efficacy of selection is S1 = N u f (N, s_1_) / u = N f (N, s_1_), where N u f (N, s_1_) is the rate of accumulation advantageous mutations. This numerator is KA (the number of nonsynonymous substitutions in cancer driver genes), while the denominator, u, is KS (the number of synonymous substitutions these genes). Let Ka and Ka’ designate the number of nonsynonymous substitutions in cancer patients and non-cancer individuals, respectively. Similarly, let Ks and Ks’ be the number of synonymous substitutions those individuals.

We now denote the lifetime risk of being diagnosed with a certain type of cancer as C_R_. Thus, KA = Ka C_R_+ Ka’ (1-C_R_) and KS=Ks C_R_ +Ks’ (1- C_R_) where Ka and Ks can be obtained from the TCGA database (https://portal.gdc.cancer.gov). The KA /KS of any cancer driver gene in a cancer tissue is R (R= Ka/Ks). Method of calculating R values of cancer tissues is described in Supporting Information. In normal tissues, Ka’/Ks’ ≈ (8) (Table 1). m = Ks/Ks’ is the ratio of the average mutation numbers in cancerous and normal tissues. We estimate m ∼ 4, based on data from cancerous and normal tissues, where Ks∼35 and Ks’∼ 8 - 9. Therefore, Eq. 2 of the main text can be written as follows:

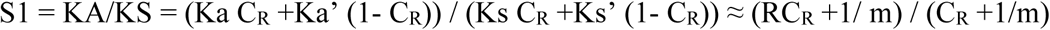

### Classification of genes according to selection intensity

COAD mutation data were downloaded from the TCGA open portal (https://portal.gdc.cancer.gov). For each data entry, TCGA implemented four mutation calling pipelines. We retained mutations called by at least two of these pipelines. To avoid super-mutated genomes, patients with more than 500 mutations in the exome were removed. The aggregated data comprise 324 COAD cases with 28788 mutations in their exomes. Normal CO and SmI mutation data are from Blokzijl et al. (21). Genes are classified in Table 1 as positively selected, neutral, or negatively selected according to each gene’s KA/KS ratio (> 4, between 4 and 0.5 and <0.5) as described in Wu et al. (8). In calculating KA and KS, the underlying mutation pattern is obtained from the data in order to give weight to different nucleotide substitutions. We analyzed the single-nucleotide substitution matrix in four-fold degenerate sites of protein-coding genes. The mutation spectrum comprised 12 basic mutation types. Mutations in CpG sites (CpG -> TpG or CpA) were also separately considered. As suggested in Wu et al. (8), genome-wide Ks value was used in lieu of the value from each individual gene which fluctuates wildly due to the small number of mutations at each locus. KA included nonsense mutations, equivalent to KAX in Wu et al. (8).

## Acknowledgement

We would like to thank Dick Hudson, Tom Nagylaki, Weiwei Zhai, Zheng Hu, Xionglei He, Pei Lin and Yongsen Ruan for discussions and advices. This work was supported by the National Science Foundation of China (31730046, 91731301, 91331202), the 985 Project (33000-18821105), the National Basic Research Program (973 Program) of China (2014CB5420,the Science Foundation for Outstanding Young Teachers in Higher Education of Guangdong (Yq2013005) and the Fundamental Research Funds for the Central Universities (16lgjc75).

